# Hunger alters approach-avoidance behaviours differently in male and female mice

**DOI:** 10.1101/2024.11.27.625796

**Authors:** Roberta G Anversa, Gemma Goldstein, Ibrahim Syed, Harry Dempsey, Amy Pearl, Xavier J Maddern, Robyn M Brown, Felicia M Reed, Andrew J Lawrence, Leigh C Walker

## Abstract

**Background:** The decision about whether to approach or avoid a reward while under threat requires balancing competing demands. Sex-specific prioritisations (e.g. mating, maternal care), or generalised prioritisations (e.g. feeding, drinking, sleeping) may differently influence approach-avoidance behaviours based on the level of “risk” and homeostatic need state of the organism. However, given known sex differences in key aspects that may influence this behaviour, direct comparison of how male and female mice make decisions to approach or avoid a dangerous area while in a fasted state have yet to be conducted.

**Methods:** We conducted several approach-avoidance tasks with varied levels of risk and reward in male and female mice that were either fasted or sated (fed). Mice underwent a light-dark box, elevated plus maze, baited large open field and runway task to assess their approach and avoidance behaviour.

**Result:** In the light-dark box and elevated plus maze, when no reward was available, fasted female mice showed greater approach behaviours than male counterparts. In the baited large open field, when reward was available, both sexes showed increased approach behaviours when fasted. However, when sated, male mice conversely showed greater approach behaviours compared to sated female mice. In the runway task, while sated mice failed to learn, fasted male mice inhibited their reward consumption in response to increased shock intensity; however, fasted female mice were resistant to increased shock intensity.

**Conclusions:** Our study identifies sex differences in decision making behaviour in mice based on satiety state across a number of approach-avoidance tasks. We highlight several nuances of these differences based on reward availability and punishment intensity. These results shine a lens on fundamental differences between the sexes in innate, survival driven behaviours that should be taken into account for future studies.

**Plain English summary:** Everyday decision making is often accompanied by conflict - whether we make the most appropriate decision or not can be influenced by both internal and external factors. Environmental threats and physiological pressures, such as hunger, can influence decision-making processes skewing the risk/reward ratio, yet how this may differ between the sexes has not been explored in detail. Here we used several tasks that assess decision-making in mice while manipulating the levels of risk or reward. Our findings show fasted female mice are more willing to engage in “risky” behaviour compared to fed female mice when risk levels were low, and no food reward was available. However, when a food reward was available, but risk levels were low, both male and female fasted mice were more likely to engage in risky behaviour compared to fed mice. Finally, when risk levels were high and food reward was available, fasted female mice continued to engage in risky behaviour, while male fasted mice were not. Together our study identifies nuanced sex differences in how male and female mice make decisions influenced by both physiological (hunger) and environmental threats and highlight the importance of understanding fundamental differences between the sexes in behaviour.

**Highlights:**

- Fasted female mice showed greater approach behaviours compared to fasted male counterparts in tasks without reward availability.
- Fasted mice of both sexes displayed greater approach behaviours when a reward was available, compared to sated controls.
- Fasted male mice inhibited reward consumption under increased shock intensity, whereas fasted female mice were resistant to mild foot shock.

## Introduction

When making decisions, both benefits and risks must be considered, requiring integration of various internal and external stimuli, and usually prioritised based on current physiological needs and/or reproductive goals of the organism (McNamara and Houston, 1986; McFarland, 1977). When an animal is hungry, it will engage in foraging behaviour; however, an approach-avoidance conflict can arise when there are two divergent goals, creating competition between incompatible motivations (Miller, 1944). For example, when foraging for food, an animal must leave its nest, which may put it at risk of predation. In this situation, an organism must dynamically calculate the value of the food reward relative to the risks associated (Gilliam and Fraser, 1987). Core aspects of decision making under this motivational conflict can be modelled using approach and avoidance behaviours. First described by Gray in the Reinforcement Sensitivity Theory, there are two main motivational systems engaged by a stimulus to achieve a goal - the behavioural approach system (move towards a reward) and the behavioural inhibition system (avoid a punishment) (Gray, 1970; Dunnet, 2004).

Rodent models of “risky” decision making often utilise a variable reward schedule of small safe or large risky reward using rodent gambling tasks. However, these models do not encapsulate the threat associated with survival, given the “risk” in these tasks is associated with delay or reward omission and a subject can chose to disengage with the task completely (Visser et al., 2011; Lockie et al., 2017). Several groups have modified classical approach-avoidance behavioural tests (light-dark box, elevated plus maze, large open field) to include a foraging element to probe how physiological hunger signals modulate decision making (Lockie et al., 2017; Teegarden and Bale, 2007). Indeed, rodents are generally more likely to engage in approach behaviours after fasting (Li et al., 2019; Towers et al., 2017; Lockie et al., 2017); however, the majority of these studies have focused on male subjects despite known sex differences in decision making (Orsini & Setlow, 2017), stress/threat sensitivity (Bangasser et al., 2019); reward seeking (Maddern et al., 2024) and metabolic hormone signalling (Börchers et al., 2022; Pearl et al., 2024; Asarian & Geary 2013). Further understanding of how biological sex may impact approach-avoidance decision making is required.

Reproductive maturity can differentially affect behavioural decision-making between sexes based on both sex-specific prioritisations (e.g. mating, maternal care), or generalised prioritisations (feeding, drinking, sleeping) (Mowrey et al., 2012). Therefore, behavioural decision-making may be highly specialised in each sex, depending on homeostatic need state. Therefore, here we assessed approach-avoidance behaviours across several approach-avoidance tasks with varying degree of reward and risk in adult male and female mice to assess how mice adapt their decision-making under conflict, dependent on energy status.

## Materials and methods

### Animals

C57BL6J mice (N = 65) were obtained from ARC (Animal Resources Centre, WA, Australia) at ∼6 weeks of age. Mice were acclimatised to a reverse light cycle (lights off 8:00-20:00 AEST). All mice had *ad libitum* access to food (laboratory chow, Barastoc, VIC, Australia) and water except where detailed. Mice were habituated to peanut butter chips (Reese’s, The Hersey Company, Hershey, PA, USA) and sucrose (8% w/v) in their cage before the tasks begun to prevent neophobia. All experiments were conducted between 9:00 – 13:00 AEST. All studies were performed in accordance with the Prevention of Cruelty to Animals Act (2004), under the guidelines of the National Health and Medical Research Council (NHMRC) Code of Practice for the Care and Use of Animals for Experimental Purposes in Australia (2013), in accordance with the ARRIVE guidelines (Percie du Sert et al., 2020) and approved by The Florey Animal Ethics Committee.

### Light-Dark Box

Male and female mice with free access to food (n = 8/sex) or which had been fasted overnight (∼16 hours; n = 8/sex) were tested in automated locomotor cells (Activity Test Chamber #ENV-510; Med Associates Inc, USA) as previously published (Maddern et al., 2024; Anversa et al., 2024). The locomotor cell was equally divided into 2 distinct zones (“light” and “dark”) connected with a small opening (10cm x 10cm) enabling mice to move freely between the zones. The “light” zone was lit by an array of light-emitting diodes (750 LUX) that were centrally placed. Mice were placed into the dark zone to begin the 10 min test, with movement tracked by Tru Scan 2.03 software (Coulbourn Instruments, USA). Mice were given at least 5 days free feeding before undergoing Elevated Plus Maze (EPM) testing.

### Elevated Plus Maze (EPM)

The elevated plus maze (EPM) apparatus was a standard mouse plus maze (30 cm [L] × 5 cm [W] × 15 cm [H]) in a room with lighting adjusted to 175 LUX with illuminance within the maze at ∼15 LUX. Fed (n = 8 males, 9 females) or fasted mice (∼16 hours fasting overnight; n = 8 males, 7 females) were placed at the junction of the four arms of the maze facing an open arm, and the duration in each arm was automatically tabulated with TopScan TopView analysing system 3.0 (CleverSyst Inc, VA, USA) for 5 min. Mice were given at least 5 days free feeding before undergoing subsequent testing.

### Baited Large Open Field (LOF)

The baited Large Open Field (LOF) was conducted as previously published (Lockie et al., 2017). The open field arena consisted of a circular area (100 cm diameter) and was illuminated with a bright light (>1000 LUX). The arena was separated into three areas, an outer zone, which extended 100 mm from the outer wall, a centre zone (10 cm diameter), in which a peanut butter chip (Reese’s™) was placed in the middle and secured to the floor of the arena with Blu-Tack (Bostic) and a transition zone between. Mice that either had access to food (fed, n = 8/sex) or were fasted overnight (∼16 hours; n = 8/sex) were placed into the transition zone and allowed to explore the arena without interruption for 10 min. Tracking software TopScan TopView analysing system 3.0 (CleverSyst Inc) automatically tracked the mice within the defined zones. After 10 min, mice were removed from the arena and placed back in the home cage with *ad libitum* access to food and water. The peanut butter chip reward was removed and weighed.

### Runway task

The runway task was conducted in a separate cohort of male (n = 18) and female (n = 15) mice as previously described (Choi et al., 2022) that were either free feeding or food restricted. Food restricted mice were maintained at 85-90% free feeding weight, starting 2 days before training. For the runway task mice were placed into a Perspex linear track (120 cm [L] x 15 cm [W] x 40 cm [H]) that was divided into 3 sections: the Start box (22 cm) constructed from grey Perspex walls and floor; the runway (88 cm) constructed of white Perspex floor and walls; and the Goal box (10 cm) constructed from grey Perspex walls and a stainless-steel grid floor. The Start box was separated from the runway with a transparent Perspex door; and the Goal box contained a stainless-steel receptacle extending 3 cm from the end wall to which the reward is delivered. The level of illumination of Start and Goal boxes was ∼200 LUX, while the runway proper was ∼350 LUX.

For training, mice underwent 4 trials per day for 4 days. Trials commenced with manual removal of the door between the Start box and the runway, with 5 µl of 8% w/v sucrose liquid/solution available from the receptacle in the Goal box on each trial. The trial was manually ended after mice consumed the sucrose reward or timed out after 120 seconds. At the end of each trial, mice were returned to the Start box for ∼30 s until the start of the next trial. Following training mice had 1 day of rest before beginning the conflict task. For conflict, fasted mice underwent the same procedure; however, the grid floor in the Goal box was electrified using a 0.05, 0.075, or 0.1 mA current. Mice received 1 day of conflict at 0.05 mA (D5), 2 days at 0.075 mA (D6-7), and 2 days at 0.1 mA (D8-9). Mice were automatically tracked using TopScan TopView analysing system 3.0 (CleverSyst Inc) within the defined zones.

### Behavioural z-score normalisation

Behavioural z-score normalisation was calculated for each animal for the following outputs from each behavioural test: Light-dark box: % time in the light side, % time in the dark side; Baited open field: % time in the target zone, % time in the outer zone; Elevated plus maze: % time in the open arms, % time in the closed arms. Given fed mice were not tested under conflict, the runway task was not integrated in the approach or avoidance score. The following formula was used to calculate each individual z-score per test using the observed output (X), population mean (µ) and population standard deviation (σ) (Labots et al., 2017; Kraeuter, 2023).

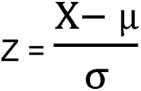

Next an integrated approach or avoidance z-score was calculated by taking the average across individual z-score from each test included in the analysis: Integrated approach score – Light-dark box: % time in the light side; Baited open field: % time in the target zone; Elevated plus maze: % time in the open arms. Integrated avoidance score – Light-dark box: % time in the dark side; Baited open field: % time in the outer zone; Elevated plus maze: % time in the closed arms.

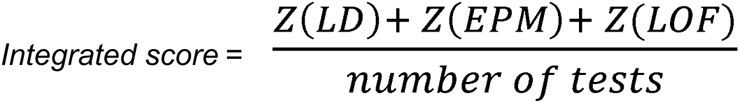

### Data analysis

Behavioural profiling was conducted in a second-by-second manner for the runway task on recorded videos from D5 (0.05 mA) and D9 (0.1 mA) using BORIS open-source software (Fuzesi et al., 2016). Approach (moving towards reward), avoidance (moving away from reward), and other behaviours (sitting, surveying, sniffing, non-directional walking and rearing) were scored. Statistical analyses were performed in GraphPad Prism version 10. Analyses for overall effects of sex and genotype were with two-way ANOVAs. Repeated measures two- or three-way ANOVA was used to analyse binned data. Bonferroni’s *post hoc* analysis was used when P < 0.1, to determine if trends were driven by a specific sex. Student’s unpaired t-test was used to analyse individual between group comparisons, where appropriate. For all tests, the level of significance was set at α = 0.05. Data represented as mean ± SEM.

## Results

### Female, but not male, mice show enhanced approach behaviours in the light-dark box after fasting

Approach-related behaviour was first assessed in the light-dark box to assess the conflict between exploration of a novel environment, and potential risk of predation (Figure 1A). Two-way ANOVA for % time spent in the light side of the light-dark box showed no main effect of sex (*F_(_*_1, 28)_ = 0.3415, *P =* 0.5636); however, a trend towards a main effect of fasting (*F*_(1, 28)_ = 2.928, *P =* 0.0981) and interaction between sex x fasting (*F*_(1, 28)_ = 3.213, *P =* 0.0838) were observed. Bonferroni’s *post hoc* analysis revealed a significant difference between fed and fasted females (*P* = 0.0391), but not males (*P* > 0.9999; Figure 1B). Time course analysis in female mice revealed a main effect of time (*F*_(3.083, 43.16)_ = 5.844, *P =* 0.0018), fasting (*F* _(1, 14)_ = 14.83, *P =* 0.0018) and time x fasting interaction (*F*_(9, 126)_ = 3.073, *P =* 0.0023). Bonferroni’s *post hoc* analysis revealed a significant difference between fed and fasted females during the 6^th^ (*P =* 0.0005), 9^th^ (*P =* 0.0031) and 10^th^ (*P =* 0.0160) minutes of the test, with a trend observed in the 7^th^ minute (*P =* 0.0991, Figure 1C). No difference was observed in male mice between fasting states when assessing across time (two-way ANOVA; fasting, *F*_(1, 14)_ = 0.001340, *P =* 0.9713); however, a main effect of time (*F*_(1.568, 21.95)_ = 11.98, *P =* 0.0007) and time x fasting interaction (*F*_(9, 126)_ = 2.148, *P =* 0.0300) were observed. No *post hoc* differences were observed between fed and fasted male mice at any timepoint (*P* > 0.05, Figure 1D). Two-way ANOVA did not report main effects of sex (*F*_(1, 28)_ = 0.1073, *P =* 0.7457), fasting (*F_(_*_1, 28)_ = 0.3052, *P =* 0.5850) or interaction (*F*_(1, 28)_ = 0.1480, *P =* 0.7033) in latency to enter the light side (Figure 1E). No difference in distance travelled was observed (two-way ANOVA; sex, *F*_(1, 28)_ = 7.4430, *P =* 0.9932; fasting, *F*_(1, 28)_ = 0.0219, *P =* 0.8832; interaction, *F*_(1, 28)_ = 1.775, *P =* 0.1936; Figure 1F).

**Figure 1.**
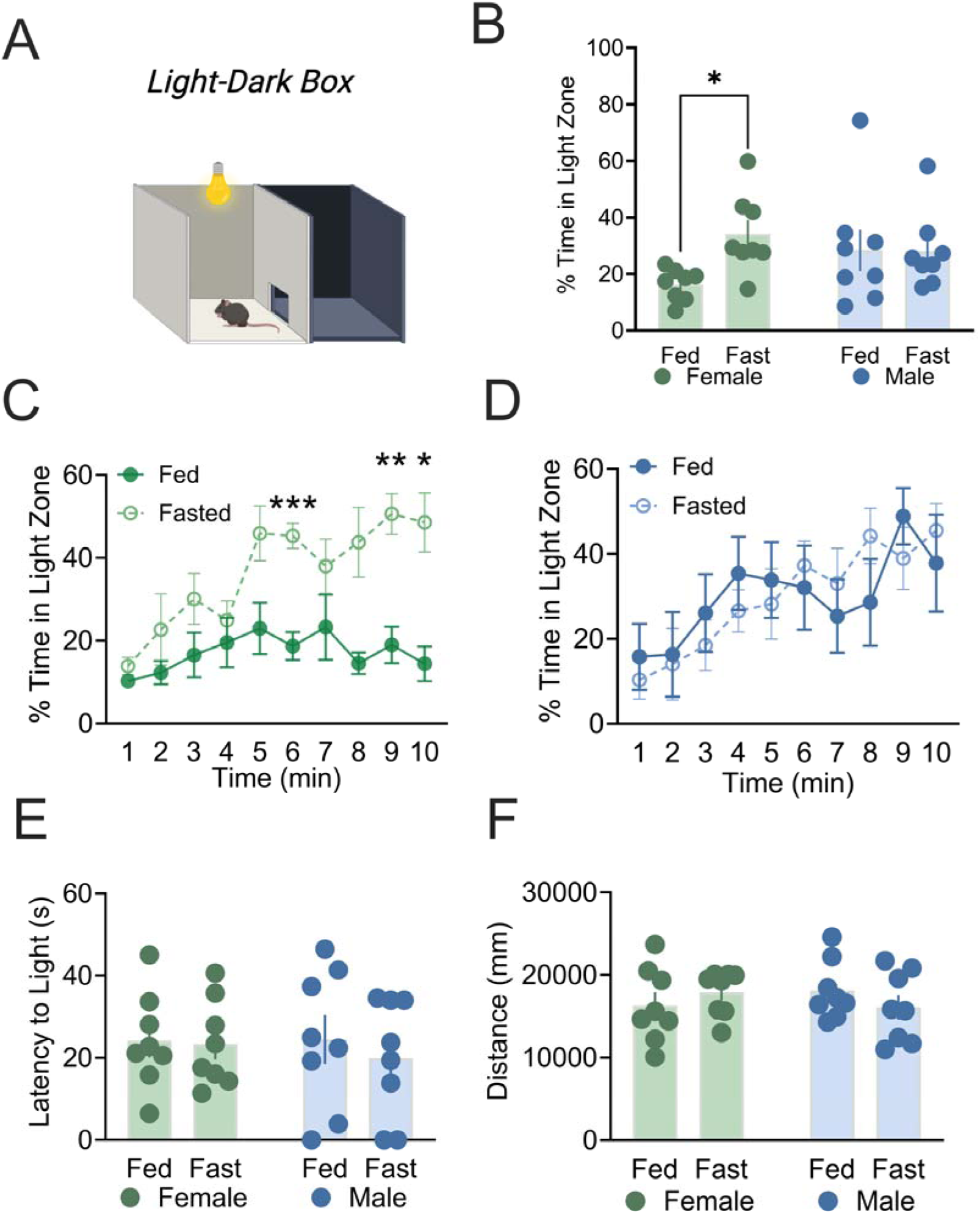
Fasting alters approach-avoidance behaviour in light-dark box of female, but not male mice. **(A)** Schematic of light-dark box. **(B)** Female fasted mice spent more time in the light zone compared to sated counterparts, while no difference was observed in males. **(C)** Differences were observed across the time course when comparing female fed vs. fasted mice. **(D)** No differences were observed between fed and fasted male mice. No difference in sex or fasting differences were observed in **(E)** latency to enter the light zone, or **(F)** distance travelled. n = 8/sex/group. *p < 0.05; **p < 0.01; ***p < 0.001. Data expressed as mean ± SEM.

### Female, but not male, mice show enhanced approach behaviours in the elevated plus maze after fasting

Approach-avoidance related behaviour was next assessed in the elevated plus maze (EPM), to assess the conflict between exploration of a novel environment, and potential risk of injury, or predation (Figure 2A). Two-way ANOVA for % time spent in the open arms of the EPM box showed no main effect of sex (*F*_(1, 28)_ = 0.1875, *P =* 0.6683), or fasting (*F*_(1, 28)_ = 2.686, *P =* 0.1124); however, a trend towards interaction between sex and fasting (*F*_(1, 28)_ = 2.965, *P =* 0.0961) was observed. Bonferroni’s *post hoc* analysis revealed a significant difference between fed and fasted females (*P* = 0.0495), but not male mice (*P* > 0.9978; Figure 2B). Time course analysis of cumulative time spent in the open arms of female mice revealed a main effect of time (*F*_(2.140, 29.96)_ = 65.37, *P* < 0.0001), a trend towards fasting (*F*_(1, 14)_ = 3.275, *P =* 0.0918) and time x fasting interaction (*F*_(4, 56)_ = 9.289, *P* < 0.0001). Bonferroni’s *post hoc* analysis revealing a significant difference between fed and fasted females during the 4^th^ (*P =* 0.0387) and 5^th^ (*P =* 0.0079) minutes of the test (Figure 2C). In male mice, a main effect was seen across time (two-way ANOVA; *F*_(1.745, 24.43)_ = 47.08, P < 0.0001); however, no difference was observed between fasting states (*F*_(1, 14)_ = 1.897, *P =* 0.1900), nor an interaction (*F*_(4, 56)_ = 1.044, *P =* 0.3930; Figure 2D). Two-way ANOVA also revealed a main effect of fasting on the latency to enter the open arms (*F*_(1, 28)_ = 4.365, *P =* 0.0459), but no effect of sex (*F_(_*_1, 28)_ = 0.03671, *P =* 0.8494), or interaction (*F*_(1, 28)_ = 1.658, *P =* 0.2085). Bonferroni’s *post hoc* analysis revealed a significant difference between fed and fasted females (*P* = 0.0483), but not male mice (*P* > 0.8183), with female fasted mice taking longer to enter the open arms (Figure 2E). No main effect of fasting (two-way ANOVA; *F*_(1, 28)_ = 0.2473, *P =* 0.6229) or sex (*F*_(1, 28)_ = 0.0454, *P =* 0.8328) nor interaction (*F*_(1, 28)_ = 2.226, *P =* 0.1469) was observed on # bouts into the open arms (Figure 2F). A trend towards an effect of fasting was observed on total distance travelled (*F*_(1, 28)_ = 3.233, *P =* 0.0830), although no main effect of sex (*F*_(1, 28)_ = 0.04906, *P =* 0.8263), or fasting x sex interaction (*F*_(1, 28)_ = 1.920, *P =* 0.1768) were observed. Bonferroni’s *post hoc* analysis revealed a trend towards decreased total distance travelled in fasted male (*P* = 0.0626), but not female mice (*P* = 0.9487; Figure 2G) when compared to their fed counterparts.

**Figure 2.**
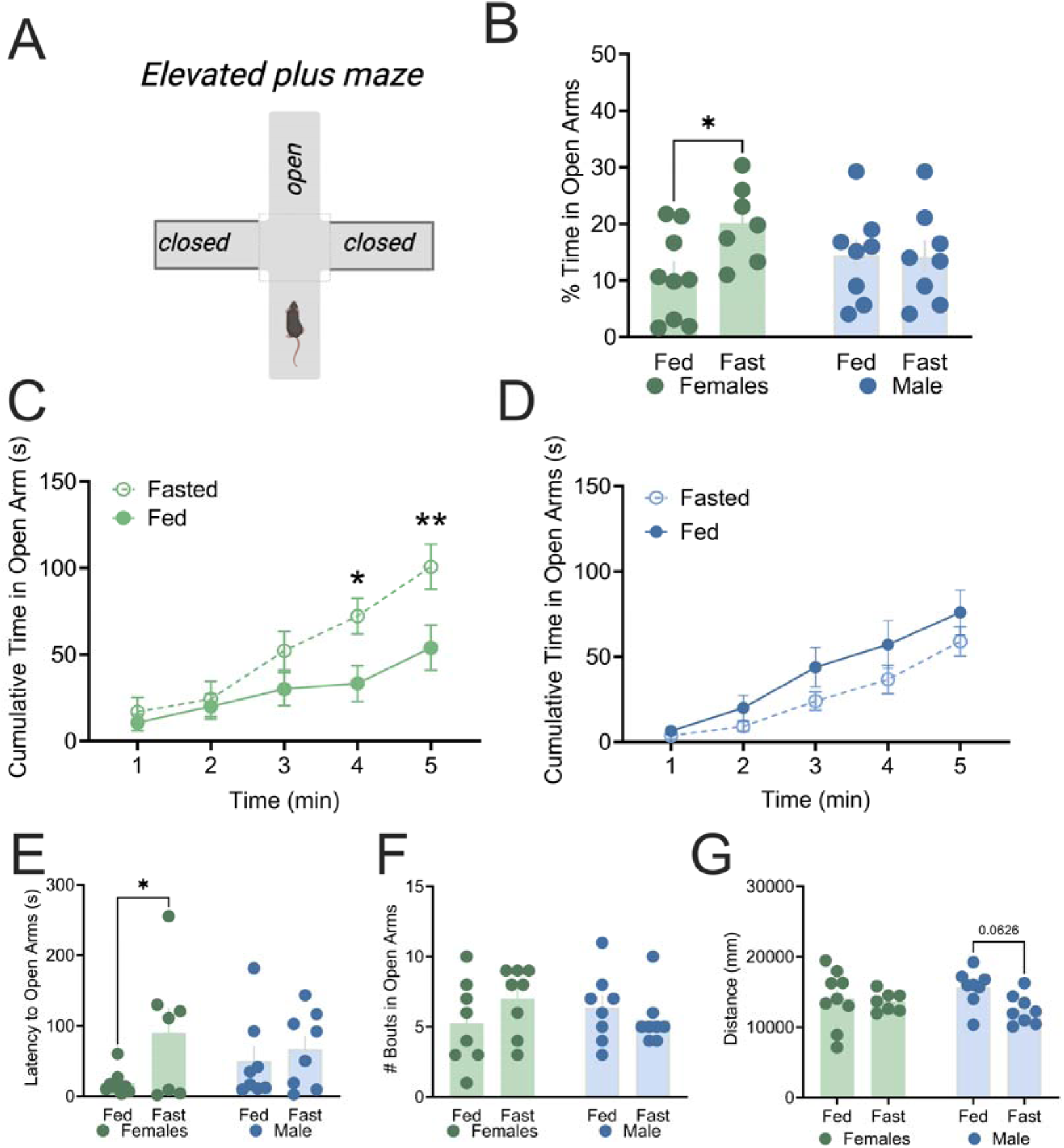
Fasting alters approach-avoidance behaviour of female, but not male mice in the elevated plus maze. **(A)** Schematic of elevated plus maze. **(B)** Female fasted mice spent more time in the open arms compared to sated counterparts, while no difference was observed in males. **(C)** Differences were observed across the time course when comparing female fed vs. fasted mice. **(D)** No differences were observed between fed and fasted male mice. **(E)** Female, but not male mice showed an increased latency to enter the open arms. No difference in sex or fasting differences were observed in **(F)** bouts in the open arms, or **(F)** distance travelled. N = 7-9/sex/group. *p < 0.05; **p < 0.01. Data expressed as mean ± SEM.

### Fasted mice of both sexes show enhanced approach behaviours in the baited large open field

Given the light-dark box and EPM tasks do not include a motivation to approach a “risky” environment, we next employed the baited large open field (LOF) (Lockie et al., 2017; Figure 3A). This task includes a peanut butter chip in the centre of the apparatus to increase conflict between reward approach or avoidance decision making behaviour. For total distance travelled, an interaction was observed between fasting and sex (*F*_(1, 27)_ = 6.031, *P =* 0.0208), with a trend towards main effect of sex (*F*_(1, 27)_ = 4.034, *P =* 0.0547), but no main effect of fasting (*F*_(1, 27)_ = 0.02566, *P =* 0.8739). Bonferroni’s *post hoc* analysis revealed a significant decrease in distance travelled by fasted male compared to fasted female mice (P= 0.0068, Figure 3B). Two-way ANOVA for amount of reward consumed showed a main effect of fasting (*F*_(1, 27)_ = 5.710, *P =* 0.0241); however, no effect of sex (*F*_(1, 27)_ = 0.1987, *P =* 0.6593), nor interaction between sex x fasting (*F*_(1, 27)_ = 0.3785, *P =* 0.5435) were observed (Figure 3C). *Post hoc* analysis indicated a trend between fed and fasted females (P = 0.0792). In line with this, a main effect of fasting (*F*_(1, 27)_ = 6.330, *P =* 0.0181), but no effect of sex (*F*_(1, 27)_ = 0.3128, *P =* 0.5806), or interaction (*F*_(1, 27)_ = 0.02476, *P =* 0.8761) was observed for % time spent in the target zone (Figure 3D). For latency to enter the target zone, two-way ANOVA showed no main effect of sex (*F*_(1, 28)_ = 0.5323, *P =* 0.4717), or fasting (*F*_(1, 28)_ = 0.008956, *P =* 0.9253); however, an interaction between sex x fasting (*F*_(1, 28)_ = 6.052, *P =* 0.0203) was observed. Bonferroni’s *post hoc* analysis revealed a trend towards reduced latency to enter the target in fasted female compared to fasted male mice (*P* = 0.0630; Figure 3E) but not compared to fed female mice (*P* > 0.05). No main effect was observed in number of entries to the target zone (two-way ANOVA, sex, *F*_(1, 28)_ = 0.06534, *P =* 0.8001; fasting, *F_(_*_1, 28)_ = 0.2981, *P =* 0.5894; interaction, *F_(_*_1, 28)_ = 1.213, *P =* 0.2802; Figure 3F). However, a significant sex x fasting interaction was observed in the number of entries in the middle (transition) zone (two-way ANOVA, sex, *F*_(1, 28)_ = 1.319, *P =* 0.2606; fasting, *F*_(1, 28)_ = 0.4664, *P =* 0.5003; interaction, *F*_(1, 28)_ = 9.270, *P =* 0.005). Bonferroni’s *post hoc* analysis revealed a significant decrease in entries in fasted compared to fed male (*P* = 0.0326), but not female mice (fasted vs. fed, *P* = 0.1819), and a significant reduction in middle zone entries in fasted males compared to fasted females (*P* = 0.0119; Figure 3G).

**Figure 3.**
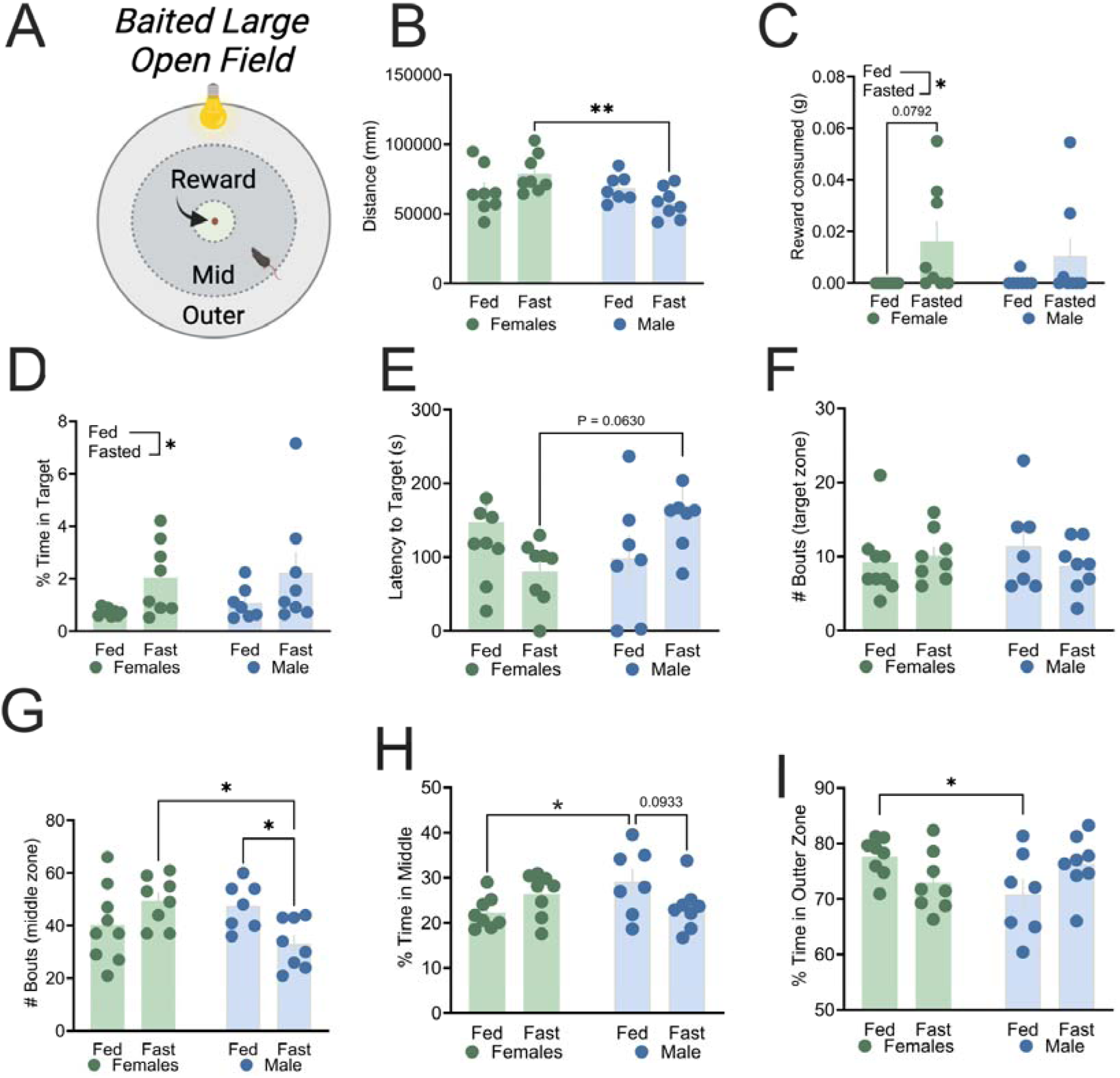
Fasting alters approach-avoidance behaviour in the elevated plus maze in female and male mice. **(A)** Schematic of baited LOF. **(B)** Distance travelled in the LOF in female (green) and male (blue) mice that were either fed or fasted prior to test. **(C)** peanut butter chip reward consumption, and **(D)** % time in target zone were increased in fasted mice. No significant difference observed in **(E)** Latency to target, or **(F)** Bout in target zone between sexes or fasting status. **(G)** Fasted male mice made less bouts into the middle (transition zone), compared to fasted females, and fed males. **(H)** Fed male mice spent less time in the middle (transition zone), compared to fed female mice’ while **(I) f**ed female mice spent more time in the outer zone, compared to fed male mice. n = 8/sex/group. *p < 0.05; **p < 0.01. Data expressed as mean ± SEM.

A significant sex x fasting interaction was also observed in the % time spent in the middle (transition) zone (two-way ANOVA, sex, *F*_(1, 27)_ = 1.028, *P =* 0.3197; fasting, *F*_(1, 27)_ = 0.1848, *P =* 0.6707; interaction, *F*_(1, 27)_ = 6.599, *P =* 0.0160). Bonferroni’s *post hoc* analysis revealed a trend towards difference between fed and fasted males (*P* = 0.0933), but not female mice (*P* = 0.2707) and a significant increase in time spent in the middle by fed male vs. fed female mice (*P* = 0.0384; Figure 3H). Further a significant sex x fasting interaction was also observed in the % time spent in the outer zone (two-way ANOVA, sex, *F*_(1, 27)_ = 0.7514, *P =* 0.3937; fasting, *F*_(1, 27)_ = 0.03956, *P =* 0.8438; interaction, *F*_(1, 27)_ = 6.641, *P =* 0.0157). Bonferroni’s *post hoc* analysis revealed a significant increase in % time spent in the outer zone for fed females, compared to fed males (*P* = 0.0477, Figure 3I).

### Female mice show enhanced integrated approach score after fasting whereas fed female mice show increased integrated avoidance score compared to male counterparts

We next calculated an integrated approach score based on time spent in the “risky” compartments on each approach-avoidance task. Two-way ANOVA revealed a significant difference in fasting (*F*_(1, 28)_ = 8.288, *P =* 0.0076), and an interaction between fasting x sex (*F*_(1, 28)_ = 10.18, *P =* 0.0035), but no effect of sex (*F*_(1, 28)_ = 1.142, *P =* 0.2943). Bonferroni’s *post hoc* analysis revealed a significant difference between fed and fasted females (*P* = 0.0011), but not male mice (*P* > 0.9999; Figure 4A). An integrated avoidance score was also calculated based on time spent in the “safe” compartments of each task. Two-way ANOVA revealed a significant difference in sex (*F*_(1, 28)_ = 5.036, *P =* 0.0329), an interaction between fasting x sex (*F*_(1, 28)_ = 4.206, *P =* 0.0498), but no effect of fasting alone (*F*_(1, 28)_ = 2.528, *P =* 0.1231). Bonferroni’s *post hoc* analysis revealed fed females show greater avoidance scores than fed males (*P* = 0.308, Figure 4B). A trend was also observed between fed and fasted females (*P* = 0.0937) and fed females compared to fasted males (*P* = 0.0697).

**Figure 4.**
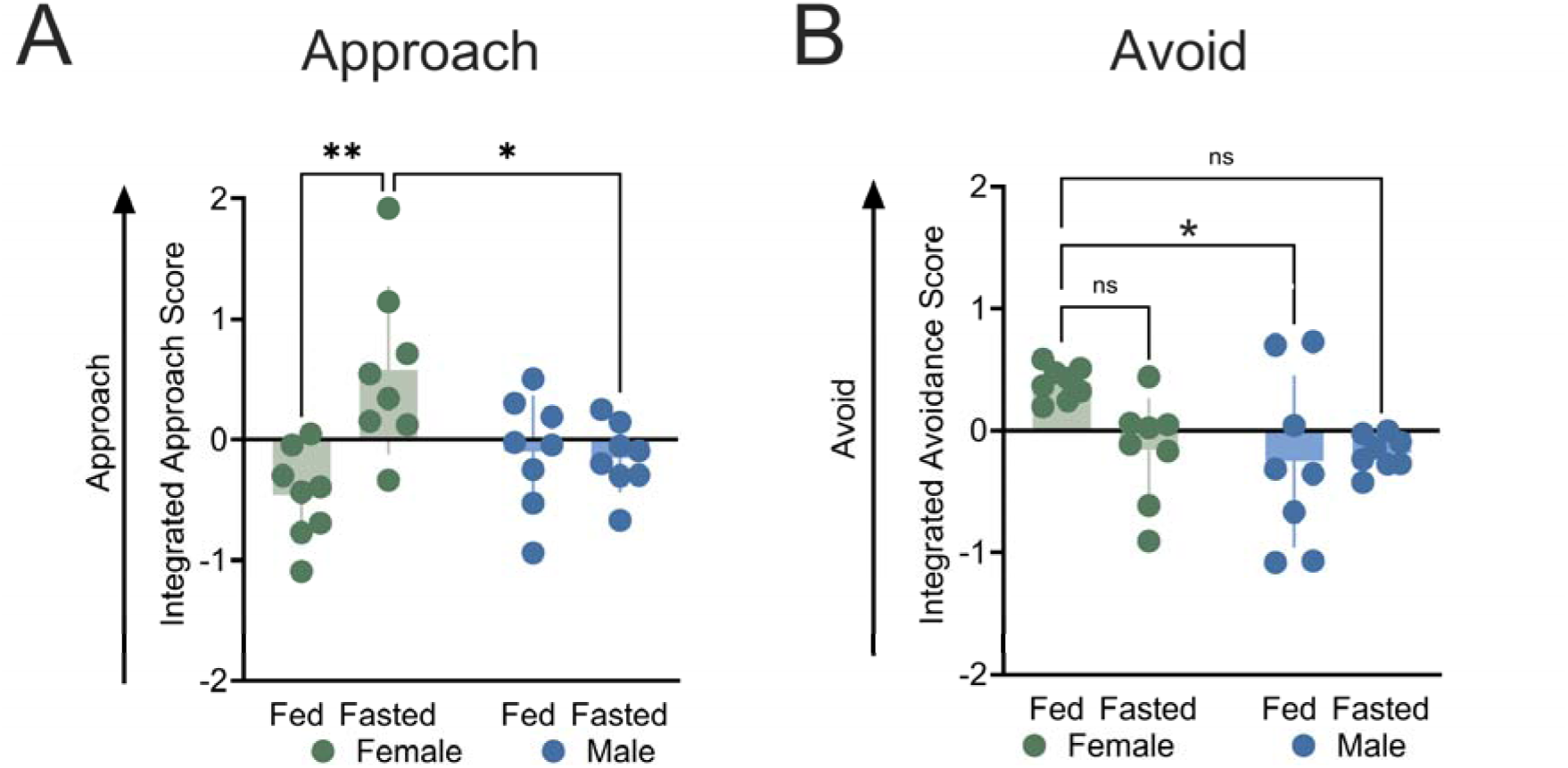
Fasting alters integrated approach and integrated-avoidance scores dependent on sex. **(A)** Female fasted mice showed an increased integrated approach score compared to fed female mice, while no difference was observed in males. **(B)** Female fed mice showed an increased avoidance score compared to male fed mice, while trends were observed between fed/fasted females and fed females with fasted males. n = 8/sex/group. *p < 0.05; **p < 0.01. Data expressed as mean ± SEM.

### Female, but not male, mice are resistant to mild punishment in the runway task after fasting

To assess approach-avoidance behaviour with the presence of a known punishment, we performed the runway task (Choi et al., 2022; Figure 5A). During training, a main effect of fasting (three-way ANOVA, *F*_(1, 29)_ = 14.15, *P* = 0.0008), a trend towards main effect of session (*F*_(3, 87)_ = 2.260, *P* = 0.0871), but no effect of sex (*F*_(1, 29)_ = 0.2273, *P* = 0.6371) or any interaction between sex, time or fasting were observed (P’s > 0.05, Figure 5B). Fed mice overall showed an average latency of 93 ± 5 (female) and 104 ± 5 (male) seconds to obtain the reward in the runway task without the introduction of conflict, compared to an average of 64 ± 3 (female) and 78 ± 11 (male) seconds for fasted mice. Given this, fed mice did not move to the conflict stage of the task.

**Figure 5.**
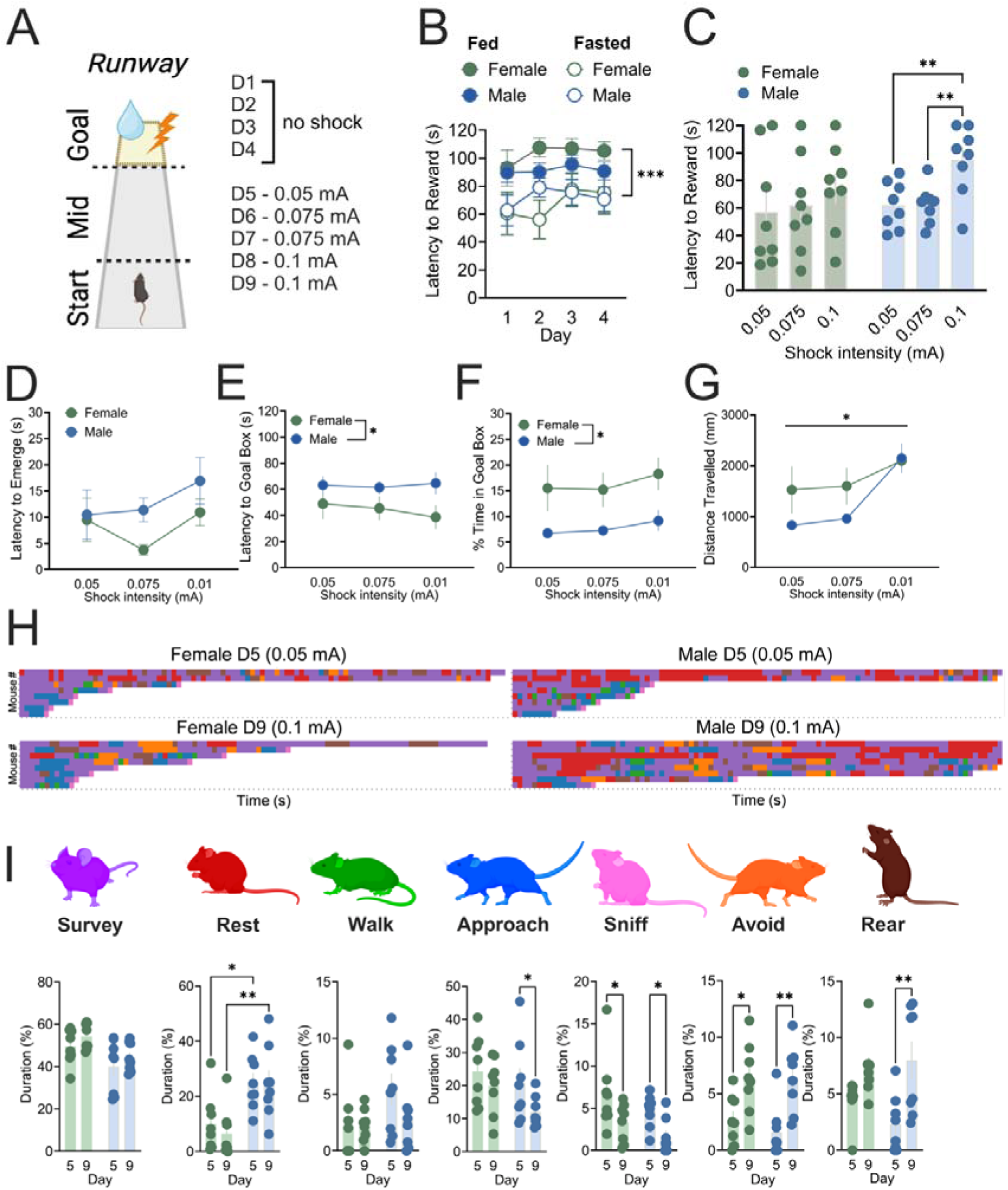
Female mice show enhanced approach behaviours in the runway task, despite punishment. (**A)** Schematic of runway task and punishment schedule. **(B)** Fasted male and female mice performed better in the training phase of the runway task. **(C)** Male, but not female mice inhibited their approach behaviour in response to a mild foot shock. **(D)** No difference observed in latency to emerge from the start box, **(E)** females showed reduced latency to enter the goal box and **(F)** spent more % time in the goal box. **(G)** Mice showed increased distance travelled in the runway with increasing shock intensity, independent of sex. **(H)** Representative behavioural barcoding map of runway trial on D5 (0.05 mA) and D9 (0.1 mA) in female (left) and male (right). Grouped analysis of duration time **(I)** Surveying (purple), resting (red), Walking (green), approaching (blue), sniffing (pink), Avoiding (orange) and rearing (brown) in mice on D5 and D9 of the runway task. n = 8-10/sex/group. *p < 0.05; **p < 0.01. Data expressed as mean ± SEM.

During conflict two-way RM ANOVA showed a main effect of shock intensity on latency to reward after conflict was introduced (F_(2, 28)_ = 10.89, *P* = 0.0003), with no effect of sex (F_(1, 14)_ = 0.4534, *P* = 0.5117), or interaction (F_(2, 28)_ = 1.625, *P* = 0.2149). Bonferroni’s *post hoc* analysis showed no significant difference between male and female mice at shock intensities (*P* > 0.05); however, a significant increase in latency to reward was observed at 0.1 mA compared to 0.05 mA (*P* = 0.0013) and 0.075 mA (*P* = 0.0016) conflict in male mice, but not female mice (*P* > 0.05; Figure 5C).

When directly comparing male and female mice, no main effect of sex (*F_(_*_1, 14)_ = 1.929, *P* = 0.1866), shock intensity (*F*_(2, 28)_ = 2.396, *P* = 0.1095) or interaction (*F_(_*_2, 28)_ = 0.6927, *P* = 0.5086) were observed in latency to emerge from the start box (Figure 5D); however, a main effect of sex (*F_(_*_1, 36)_ = 7.598, *P* = 0.0091), but not shock intensity (*F*_(2, 36)_ = 0.1407, *P* = 0.8692) or interaction (*F_(_*_2, 36)_ = 0.2890, *P* = 0.7507) were observed in the latency to enter the goal box (Figure 5E). A main effect of sex (*F*_(1, 14)_ = 5.644, *P* = 0.0323), but not shock intensity (*F*_(1.607, 22.49)_ = 2.042, *P* = 0.1597) or interaction (*F*_(2, 28)_ = 0.07935, *P* = 0.9239) was observed in time spent in the goal box (Figure 5F). For distance travelled, a main effect of shock intensity (*F*_(1.327, 18.57)_ = 12.23, *P* = 0.0013), but no effect of sex (*F*_(1, 14)_ = 1.868, *P* = 0.1933), nor interaction (*F*_(2, 28)_ = 1.960, *P* = 0.1597) were observed (Figure 5G).

Next, we compared behavioural profiles across the first day of conflict (D5; 0.05 mA intensity) and the fifth day of conflict (D9; 0.1 mA) (Figure 5H-I). RM two-way ANOVA showed a main effect of sex (*F*_(1, 14)_ = 10.27, *P* = 0.0064), but not conflict intensity on % time spent surveying (*F*_(1, 14)_ = 1.941, *P* = 0.1853), with Bonferroni’s *post hoc* analysis showing no specific differences between days (*P* > 0.05) (Figure 5I). A significant main effect of sex was observed on % time spent resting (*F*_(1, 14)_ = 15.28, *P* = 0.0016), with Bonferroni’s *post hoc* analysis showing this to be significantly more in males on D1 (*P* = 0.0189) and D5 (*P* = 0.0040) compared to females (Figure 5I). No significant difference in non-directional walking was observed (shock intensity, *F*_(1, 14)_ = 1.615, *P* = 0.2245; sex, *F*_(1, 14)_ = 2.826, *P* = 0.1149; interaction, *F_(_*_1, 14)_ = 1.609, *P* = 0.2254). A main effect of shock intensity was observed on approach behaviours (F_(1, 14)_ = 7.624, *P* = 0.0153), with no main effect of sex (*F*_(1, 14)_ = 1.904, *P* = 0.1893) or interaction (*F*_(1, 14)_ = 0.9754, *P* = 0.3401). Bonferroni’s *post hoc* analysis showing a significant decrease in % time approaching (walking towards reward) in male (*P* = 0.0380), but not female mice (*P* = 0.4607; Figure 5I). A significant effect of shock intensity was observed in % time spent sniffing food (*F*_(1, 14)_ = 19.23, *P* = 0.0006). Bonferroni’s *post hoc* analysis showed a significant decrease in sniffing behaviour in both male (*P* = 0.0219), and female (*P* = 0.0112) mice between D5 and D9 (Figure 5I). Further, a main effect of shock intensity was observed in % time avoiding the reward (*F*_(1, 14)_ = 20.67, *P* = 0.0005). Finally, a significant main effect of shock intensity was observed in rearing behaviour (*F*_(1, 14)_ = 14.67, *P* = 0.0018). Bonferroni’s *post hoc* analysis showed this increase was driven by male (*P* = 0.0061), but not female mice (P = 0.1744, Figure 5I).

## Discussion

The decision about whether to approach or avoid a reward while under threat requires balancing competing demands. Sex-specific prioritisations (e.g. mating, maternal care), or generalised prioritisations (feeding, drinking, sleeping), may differently influence approach-avoidance behaviours based on the level of “risk” and homeostatic need state of the organism. Therefore, we investigated how both levels of “risk”, and satiety (fed vs fasted) alter approach-avoidance behaviour in male and female mice. Overall, we highlight novel sex differences in how fasting alters approach-avoidance decision-making that depend on reward availability and level of risk in the task. Specifically, we show female, but not male, mice increase approach behaviours in the light-dark box and EPM after fasting, without food available. However, in the baited LOF, where a food reward is available, both sexes showed enhanced approach behaviour. Finally, neither male nor female mice were able to learn the runway task when sated; however, when fasted, female mice were more resistant to the mild foot shock, showing increased approach behaviours under high risk probability and reward availability.

To assess approach-avoidance conflict, our mice were subjected to distinct behavioural tasks that varied in the availability of reward, and the level of risk. In the light-dark box and EPM, which have relatively low risk and no food reward availability, fasted female mice showed greater approach/exploratory behaviours compared to fed counterparts, while no difference was observed in fed vs. fasted males. Previous studies have shown no difference in how male mice explored the light-side of the light-dark box after overnight (∼16h) fasting (Lockie et al., 2017); however, an increase in time spent in the open arms of the EPM was observed in both sexes after 24 h fasting in another study, although sexes were combined and sex effects not reported (Li et al., 2019). Interestingly, this effect differed based on circadian variation, with differences observed in the early, but not late light period; and 12 h fasting was not sufficient to increase time in the open area of the open field task (Li et al., 2019). Of note, the current experiments were conducted in the early-mid dark phase, when mice would naturally engage in foraging behaviours (Kronfeld-Schor and Dayan, 2008), and after overnight (∼16 hours) fasting, which may explain some differences. Seasonal variations in activity and age-related variations have also been reported (Pernold et al., 2021). Our experiments were all undertaken in early adult mice (8-12 weeks ages throughout) and in the summer/autumn of the southern hemisphere in matched cohorts, but the impact of these variations cannot be ruled out. Collectively, these data suggest that fasting state is a major factor in decision-making under conflict in females, who are more willing to explore “risky” areas, despite no food availability.

To assess the generalisability of sex differences in approach-avoidance after fasting we increased the motivational conflict by employing a baited LOF (Lockie et al., 2017). Here our findings that fasted male mice had increased motivational drive and consummatory behaviour compared with sated male mice, align with previous studies (Lockie et al., 2017; Dodd et al., 2021). We extend these findings to show similar behaviour in fasted female mice. Interestingly, while approach and consummatory behaviours were similar between the sexes, where they decided to spend their time outside of the target (reward) zone showed distinct differences between the sexes, based on fasting state. When fed, male mice were more likely to spend time in the middle (transition) zone and, conversely, fed female mice spent more time in the outer (safe) zone. This aligns with our integrated avoidance score, which showed fed females spend less time in the “risky” zones compared to fed males, and previous literature suggesting female mice are generally more risk averse than male counterparts (Francesconi et al, 2020; Orsini et al., 2016). To our knowledge this is the first assessment of female mice risk-taking behaviour using this task and highlights nuances in decision making between the sexes based on ability to obtain a reward. Independent of reward availability, fasted female mice were more likely to explore risky areas; however, when sated, they showed less risky behaviour compared to males.

To determine how intensity of “risk” may impact decision making between the sexes we employed the runway task (Choi et al., 2022). Here, after training, mice must overcome an electrified grid to receive a reward. Our training showed that fed mice did not consistently collect the reward, even in the absence of punishment, therefore we only continued to the conflict stage of the task with fasted mice. During conflict, fasted female mice showed greater approach behaviours, which were resistant to increases in shock intensity. As shock intensity was increased, male mice inhibited their behaviour, consistent with previous studies (Choi et al., 2022). When examining behavioural phenotypes male mice spent more % time resting compared to females, aligning with previous literature suggesting female rodents show greater locomotor activity under similar conditions (Maren et al., 1994; Aguilar et al., 2003; Bethancourt et al., 2011). Increased shock intensity enhanced avoidance behaviour (moving away from reward) and sniffing the reward in both sexes, while rearing was increased, and approach behaviour reduced specifically in males during high conflict compared to females. Previous studies have shown rearing increases after stress in male mice (Füzesi et al., 2016), suggesting the task may elicit a stress response in fasted males; however, this was not the case in female mice. Sex differences in response to a foot shock have been previously described, with female rodents freezing less than male counterparts (Dalla & Shors, 2009; Aguilar et al., 2003; Bethancourt et al., 2011). Studies suggest this may be driven by different types of fear responses, not pain thresholds, with females showing greater levels of active fear response (darting) behaviours than males (Gruene et al., 2015; Dalla & Shors, 2009). Therefore, sex differences observed in the runway task may be driven by active vs. inactive responses to the electrified grid during conflict.

Another factor that may contribute to the sex differences observed is learning. Previous research using an explore/exploit task, the spatial restless two-armed bandit, reported that male mice had higher levels of exploration than females throughout the task, but that was due to males being more likely to get ‘stuck’ exploring before committing to a favoured choice. In contrast, female mice showed elevated reward-driven learning specifically during shorter bouts of exploration, which led them to start exploiting a favoured choice earlier than males (Chen et al., 2021). Other studies have shown sex differences in stress-induced binge eating, with fed female, but not male mice displaying binge-like behaviour after a mild psychological stress (Anversa et al., 2020; 2023). Further, differences between the sexes may be driven by changes in demand for highly palatable foods (Freeman et al., 2021; Tapia et al., 2019); however, we have not observed differences in taste preferences between the sexes previously (Maddern et al., 2024).

Metabolic differences have also been reported between sexes. For example, sex differences in the levels of the orexigenic hormone ghrelin, which increases when the stomach is empty in preparation for a meal (Cummings et al., 2001) have been observed across species, with females exhibiting higher levels of acylated ghrelin in a fasted state than males (Makovey et al., 2007; Beasley et al., 2009; Tai et al., 2009; Leone et al., 2022; Borchers et al., 2022). Further, while ghrelin increases activity of midbrain neurons in both sexes, females show greater response to bath application of ghrelin (Pearl et al., 2024), and greater ghrelin receptor (GHSR) expression has been observed in the female rodent brain, including the amygdala, hippocampus, lateral hypothalamus and Edinger-Westphal nucleus, (Pearl et al., 2024; Borchers et al., 2022). Therefore, the intensity of “hunger” may be perceived differently between the sexes and/or ghrelin may exert different actions within the brain driving changes in how approach-avoidance decisions are made. However, further studies are required to understand these interactions and how the brain integrates signals of threat and hunger to make appropriate decisions.

A limitation of our study lies in the lack of oestrus cycle tracking. However, the findings were relatively consistent across tasks that were performed 3-5 days apart in the same animals (L/D box, EPM, LOF). Given the oestrus cycle in female mice can vary in duration (∼5-8 days) and be irregular between littermates (Ryherd et al., 2023), this suggests stage of the oestrus cycle is not likely a major contributor to enhanced approach behaviour in fasted female mice, although this cannot be ruled out and requires further examination. However, it should be noted that fluctuating hormone levels during adulthood are only one small aspect of biological sex differences. Organisational effects of both sex chromosome complement, epigenetic regulators, and sex hormone levels during development all contribute to fundamental sex differences that may underpin differences observed in behaviour (Maddern et al., 2024; Kopsida et al., 2013; Williams and Meck, 1991). Studies in this area are critical to determine organisational vs. activation effects of hormones and their impact on behaviour generally.

## Conclusions

In summary, here we identify sex differences in decision making behaviour in mice based on satiety state across a number of approach-avoidance tasks. We highlight several nuances of these differences based on reward availability and punishment intensity. In sated conditions male mice show greater approach behaviour compared to female mice, while when fasted, female mice have greater approach behaviours. Further in female mice, this is less sensitive to punishment, suggesting homeostatic need state drives greater foraging and reward driven behaviours in female mice. These results shine a lens on fundamental differences between the sexes in innate, survival driven behaviours that should be taken into account for future studies. Research exploring fundamental sex differences in approach-avoidance decision making, the impact across the life span, and the neurocircuitry that drive these behaviours should be investigated.

## Author contributions

Conceptualisation: RGA, FMR, AJL, LCW; Methodology: RGA, GG, HD, FMR, LCW; Data collection: RGA, GG, IS, AP, XJM; Data analysis: RGA, GG, IS, HD, LCW; Interpretation of finding and drafting manuscript: RGA, RMB, FMR, AJL, LCW; Funding acquisition: AJL, LCW. All authors have read and approved the final version of the manuscript.

## Data availability

Data are available from the corresponding author upon reasonable request.

## Declarations Ethics

No human participants were used in the current study. All studies were performed in accordance with the Prevention of Cruelty to Animals Act (2004), under the guidelines of the National Health and Medical Research Council (NHMRC) Council Code of Practice for the Care and Use of Animals for Experimental Purposes in Australia (2013), in accordance with the ARRIVE guidelines and approved by The Florey Animal Ethics Committee.

## Consent

Not applicable

## References

Aguilar R, Gil L, Gray JA, Driscoll P, Flint J, Dawson GR, et al. Fearfulness and sex in F2 Roman rats: males display more fear though both sexes share the same fearfulness traits. Physiol Behav. 2003;78(4-5):723–32.

Anversa RG, Campbell EJ, Ch’ng SS, Gogos A, Lawrence AJ, Brown RM. A model of emotional stress-induced binge eating in female mice with no history of food restriction. Genes Brain Behav. 2020;19(3):e12613.

Anversa RG, Campbell EJ, Walker LC, S SCn, Muthmainah M, F SK, et al. A paraventricular thalamus to insular cortex glutamatergic projection gates "emotional" stress-induced binge eating in females. Neuropsychopharmacology. 2023;48(13):1931–40.

Asarian L, Geary N. Sex differences in the physiology of eating. Am J Physiol Regul Integr Comp Physiol. 2013;305(11):R1215–67.

Bangasser DA, Eck SR, Ordones Sanchez E. Sex differences in stress reactivity in arousal and attention systems. Neuropsychopharmacology. 2019;44(1):129–39.

Beasley JM, Ange BA, Anderson CA, Miller Iii ER, Holbrook JT, Appel LJ. Characteristics associated with fasting appetite hormones (obestatin, ghrelin, and leptin). Obesity (Silver Spring). 2009;17(2):349–54.

Bethancourt JA, Vasquez CE, Britton GB. Sex-dependent effects of long-term oral methylphenidate treatment on spontaneous and learned fear behaviours. Neurosci Lett. 2011;496(1):30–4.

Borchers S, Krieger JP, Maric I, Carl J, Abraham M, Longo F, et al. From an Empty Stomach to Anxiolysis: Molecular and Behavioural Assessment of Sex Differences in the Ghrelin Axis of Rats. Front Endocrinol (Lausanne). 2022;13:901669.

Chen CS, Knep E, Han A, Ebitz RB, Grissom NM. Sex differences in learning from exploration. Elife. 2021;10.

Choi EA, Husic M, Millan EZ, Gilchrist S, Power JM, Jean-Richard Dit Bressel P, et al. A Corticothalamic Circuit Trades off Speed for Safety during Decision-Making under Motivational Conflict. J Neurosci. 2022;42(16):3473–83.

Cummings DE, Purnell JQ, Frayo RS, Schmidova K, Wisse BE, Weigle DS. A preprandial rise in plasma ghrelin levels suggests a role in meal initiation in humans. Diabetes. 2001;50(8):1714-9.

Dalla C, Shors TJ. Sex differences in learning processes of classical and operant conditioning. Physiol Behav. 2009;97(2):229–38.

de Visser L, Homberg JR, Mitsogiannis M, Zeeb FD, Rivalan M, Fitoussi A, et al. Rodent versions of the iowa gambling task: opportunities and challenges for the understanding of decision-making. Front Neurosci. 2011;5:109.

Dodd GT, Kim SJ, Mequinion M, Xirouchaki CE, Bruning JC, Andrews ZB, et al. Insulin signaling in AgRP neurons regulates meal size to limit glucose excursions and insulin resistance. Sci Adv. 2021;7(9).

Dunnett SB. The neuropsychology of anxiety: An enquiry into the functions of the septo-hippocampal system, 2nd edition. Psychologist. 2004;17(3):149-.

Francesconi JA, Macaroy C, Sawant S, Hamrick H, Wahab S, Klein I, et al. Sexually dimorphic behavioural and neural responses to a predator scent. Behav Brain Res. 2020;382:112467.

Freeman LR, Bentzley BS, James MH, Aston-Jones G. Sex Differences in Demand for Highly Palatable Foods: Role of the Orexin System. Int J Neuropsychopharmacol. 2021;24(1):54–63.

Fuzesi T, Daviu N, Wamsteeker Cusulin JI, Bonin RP, Bains JS. Hypothalamic CRH neurons orchestrate complex behaviours after stress. Nat Commun. 2016;7:11937.

Gilliam JF, Fraser DF. Habitat Selection Under Predation Hazard: Test of a Model with Foraging Minnows. Ecology. 1987;68(6):1856–62.

Gray JA. The psychophysiological basis of introversion-extraversion. Behav Res Ther. 1970;8(3):249–66.

Gruene TM, Flick K, Stefano A, Shea SD, Shansky RM. Sexually divergent expression of active and passive conditioned fear responses in rats. Elife. 2015;4:e11352.

Kopsida E, Lynn PM, Humby T, Wilkinson LS, Davies W. Dissociable effects of Sry and sex chromosome complement on activity, feeding and anxiety-related behaviours in mice. PLoS One. 2013;8(8):e73699.

Kronfeld-Schor N, and Dayan T. Activity patterns of rodents: the physiological ecology of biological rhythms. Biol. Rhytm. Res. 2008;9(3):193–211.

Leone A, De Amicis R, Pellizzari M, Bertoli S, Ravella S, Battezzati A. Appetite ratings and ghrelin concentrations in young adults after administration of a balanced meal. Does sex matter?. Biology of sex Differences. 2022;13(1):25.

Li C, Hou Y, Zhang J, Sui G, Du X, Licinio J, et al. AGRP neurons modulate fasting-induced anxiolytic effects. Transl Psychiatry. 2019;9(1):111.

Lockie SH, McAuley CV, Rawlinson S, Guiney N, Andrews ZB. Food Seeking in a Risky Environment: A Method for Evaluating Risk and Reward Value in Food Seeking and Consumption in Mice. Front Neurosci. 2017;11:24.

Maddern XJ, Lawrence AJ, Campbell EJ. Electric barrier-induced voluntary abstinence reduces alcohol seeking in male, but not female, iP rats. Behav Neurosci. 2024;138(1):1–14.

Makovey J, Naganathan V, Seibel M, Sambrook P. Gender differences in plasma ghrelin and its relations to body composition and bone - an opposite-sex twin study. Clin Endocrinol (Oxf). 2007;66(4):530–7.

Maren S, De Oca B, Fanselow MS. Sex differences in hippocampal long-term potentiation (LTP) and Pavlovian fear conditioning in rats: positive correlation between LTP and contextual learning. Brain Res. 1994;661(1-2):25–34.

Mcfarland DJ. Decision-Making in Animals. Nature. 1977;269(5623):15–21.

Mcnamara JM, Houston AI. The Common Currency for Behavioural Decisions. Am Nat. 1986;127(3):358–78.

Miller NE. Experimental studies of conflict. In J. M. Hunt, Personality and the behaviour disorders (pp. 431–465).1994, Ronald Press.

Mowrey WR, Portman DS. Sex differences in behavioural decision-making and the modulation of shared neural circuits. Biol Sex Differ. 2012;3:8.

Orsini CA, Setlow B. Sex differences in animal models of decision making. J Neurosci Res. 2017;95(1-2):260–9.

Orsini CA, Willis ML, Gilbert RJ, Bizon JL, Setlow B. Sex differences in a rat model of risky decision making. Behav Neurosci. 2016;130(1):50–61.

Pearl A, Pinares-Garcia P, Shesham A, Maddern X, Anversa RG, Brown RM, et al. Edinger-Westphal ghrelin receptor signalling regulates binge alcohol consumption in a sex specific manner. bioRxiv. 2024:2024.03.23.586439.

Percie du Sert N, Hurst V, Ahluwalia A, Alam S, Avey MT, Baker M, … & Würbel, H. The ARRIVE guidelines 2.0: Updated guidelines for reporting animal research. JCBFM 2020;40(9):1769–1777.

Pernold K, Rullman E, Ulfhake B. Major oscillations in spontaneous home-cage activity in C57BL/6 mice housed under constant conditions. Sci Rep. 2021;11(1):4961.

Ryherd GL, Bunce AL, Edwards HA, Baumgartner NE, Lucas EK. Sex differences in avoidance behaviour and cued threat memory dynamics in mice: Interactions between estrous cycle and genetic background. Horm Behav. 2023;156:105439.

Tai K, Visvanathan R, Hammond AJ, Wishart JM, Horowitz M, Chapman IM. Fasting ghrelin is related to skeletal muscle mass in healthy adults. Eur J Nutr. 2009;48(3):176–83.

Tapia MA, Lee JR, Weise VN, Tamasi AM, Will MJ. Sex differences in hedonic and homeostatic aspects of palatable food motivation. Behav Brain Res. 2019;359:396–400.

Teegarden SL, Bale TL. Decreases in dietary preference produce increased emotionality and risk for dietary relapse. Biol Psychiatry. 2007;61(9):1021–9.

Towers AE, Oelschlager ML, Patel J, Gainey SJ, McCusker RH, Freund GG. Acute fasting inhibits central caspase-1 activity reducing anxiety-like behaviour and increasing novel object and object location recognition. Metabolism. 2017;71:70–82.

Williams CL, Meck WH. The organizational effects of gonadal steroids on sexually dimorphic spatial ability. Psychoneuroendocrinology. 1991;16(1-3):155–76.

